# A spatial memory signal shows that the parietal cortex has access to a craniotopic representation of space

**DOI:** 10.1101/173245

**Authors:** Mulugeta Semework, Sara C. Steenrod, Michael E. Goldberg

## Abstract

Humans effortlessly establish a gist-like memory of their environment whenever they enter a new place. They can then use this memory to guide action even in the absence of vision. Neurons in the lateral intraparietal area (LIP) of the monkey exhibit a form of this environmental memory, responding when a monkey makes a saccade that brings the spatial location of a stimulus that appeared on a number of prior trials, but not on the present trial into their receptive fields. The stimulus need never have appeared in the receptive field of the neuron. This response is usually weaker with a longer latency than the neuron’s visual response. We suggest that these results demonstrate that LIP has access to a craniotopic memory of space, which is activated when the spatial location of the vanished stimulus can be described by a retinotopic vector from the center of gaze to the stimulus.

---

Humans, and presumably monkeys, effortlessly establish a gist-like memory of their environment whenever they enter a new place. They can then use this memory to guide action even in the absence of vision. The hallmark of this environmental memory is that the objects remembered need not be relevant to the subject’s current behavior. For example, you may never be asked to point to the door with your eyes closed, even though you establish the memory of the door’s location automatically. Although normally humans have no trouble pointing to objects in the room with their eyes closed, patients with parietal lesions cannot do this even though they can easily locate and point to objects in the room with their eyes open (Farah et al., 1988), suggesting an environmental memory impairment.

Visually responsive neurons in the frontal eye field (FEF) exhibit a signal that could represent environmental memory (Umeno and Goldberg, 2001). After monkeys make a number of saccades that bring a task-irrelevant probe stimulus into the receptive field of a visually responsive FEF neuron, many neurons respond on trials when a saccade brings the spatial location of the stimulus into the receptive field, even though the probe stimulus did not appear on the current trial. However, because the investigators always established the memory response by using the same saccade as they did to evoke and evaluate the memory response, it is not clear if the effect is a true spatial memory or a merely a memory of receptive field stimulation by a saccade.

Here we asked if neurons in the lateral intraparietal area (LIP), an area with visual, oculomotor, and mnemonic connections that serves as a priority map of the environment (Bisley and Goldberg, 2010), also exhibits an environmental memory signal. Indeed, we found that neurons in LIP did exhibit an environmental memory signal. Further, we found that the memory signal could be established even when the stimulus never appeared in the receptive field of the neuron, and occurred when the monkey made any saccade that brought the spatial location of the vanished probe stimulus into the receptive field. In addition, the memory response occurred even if the monkey were planning a subsequent saccade that removed the spatial location of the vanished stimulus from the receptive field. These results suggest that LIP has access to a representation of the visual world in at least craniotopic coordinates. Because saccades and reaching movements are coded in the parietal cortex in retinotopic coordinates (Duhamel et al., 1992, Andersen et al., 1998), these results suggest that one role of LIP is to transform a craniotopic representation of the remembered environment into a retinotopic representation more useful for the generation of action.

## Results

### Dataset

We recorded the activity of 288 neurons in 3 monkeys: 255 in Monkey A, 12 in monkey B, and 21 in Monkey C. Since the results from monkey B and C were comparable and we had only a small number of cells in each monkey, we combined them into “monkey B”. All neurons in our sample had visual responses to the onset of a saccade target in their receptive fields, and exhibited delay-period and/or presaccadic activity in the memory-guided delayed saccade task. Furthermore, their postsaccadic responses were not contaminated by the phasic and tonic components of an eye position signal often found in parietal neurons (Andersen et al., 1988). Finally, they exhibited no perisaccadic activity when the monkey made the saccade that would subsequently be used to evaluate the memory response. All neurons were situated in the lateral bank of the intraparietal sulcus, as determined by structural MRI (Figure 1a). We used three different tasks based on single and double-step saccade eye-movement paradigms (Figure 1b) designed to demonstrate environmental memory in LIP neurons.

**Figure 1.**
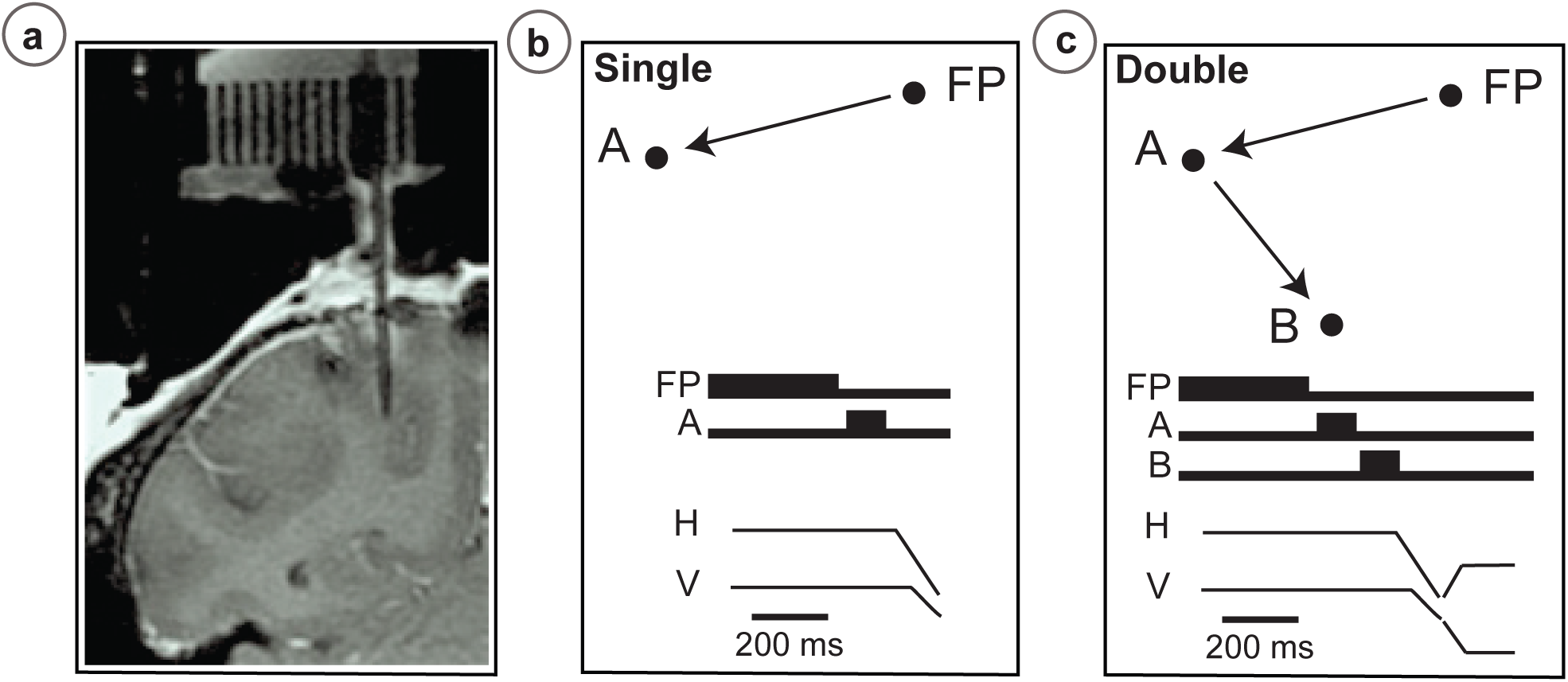
A Location of LIP recordings. A tungsten microelectrode (250 μm thick, straight shadow) located in the target area based on known LIP activity and the commonly used atlas-defined (Paxinos et al., 1999., Saleem and Logothetis, 2012) coordinates within the given coronal slice of the brain. This electrode location was at -2 AP and +10 ML. *Single (B) & double* - *step (C) saccades*. Cartoons show spatial location of fixation point and targets. Thick traces indicate the onset and duration of the FP, A and B stimuli. H and V are horizontal & vertical eye movements.

#### Task 1 - Basic Memory Task

We studied 166 LIP neurons using a task based on that used to demonstrate environmental memory in the frontal eye field (Umeno and Goldberg, 2001) (Figure 2). After determining the spatial tuning properties of the neuron being recorded, the monkeys performed the Basic Memory Task which is comprised of four blocks. The arrangement of the stimuli (fixation point, saccade target, and probe stimulus) was customized according to the spatial properties of each neuron’s receptive field. In the first block, we asked the monkey to perform a block of 20 visually-guided saccade trials in which no probe stimuli appeared on the screen (Figure 2, Block 1). In these trials no stimulus, including the saccade target, encroached on the receptive field of the neuron. Next, we asked the monkey to perform a block of 30 visually-guided saccade trials in which a task-irrelevant probe stimulus appeared at a location that would be brought into the receptive field by the required saccade (i.e., the future receptive field of the neuron) (Figure 2, Block 2). We then pseudorandomly interleaved trials in which the probe stimulus appeared on the screen with those in which no probe stimulus appeared (Figure 2, Block 3). Finally, we presented the monkey with a block of 100 trials in which, again, the probe stimulus never appeared (Figure 2, Block 4).

**Figure 2.**
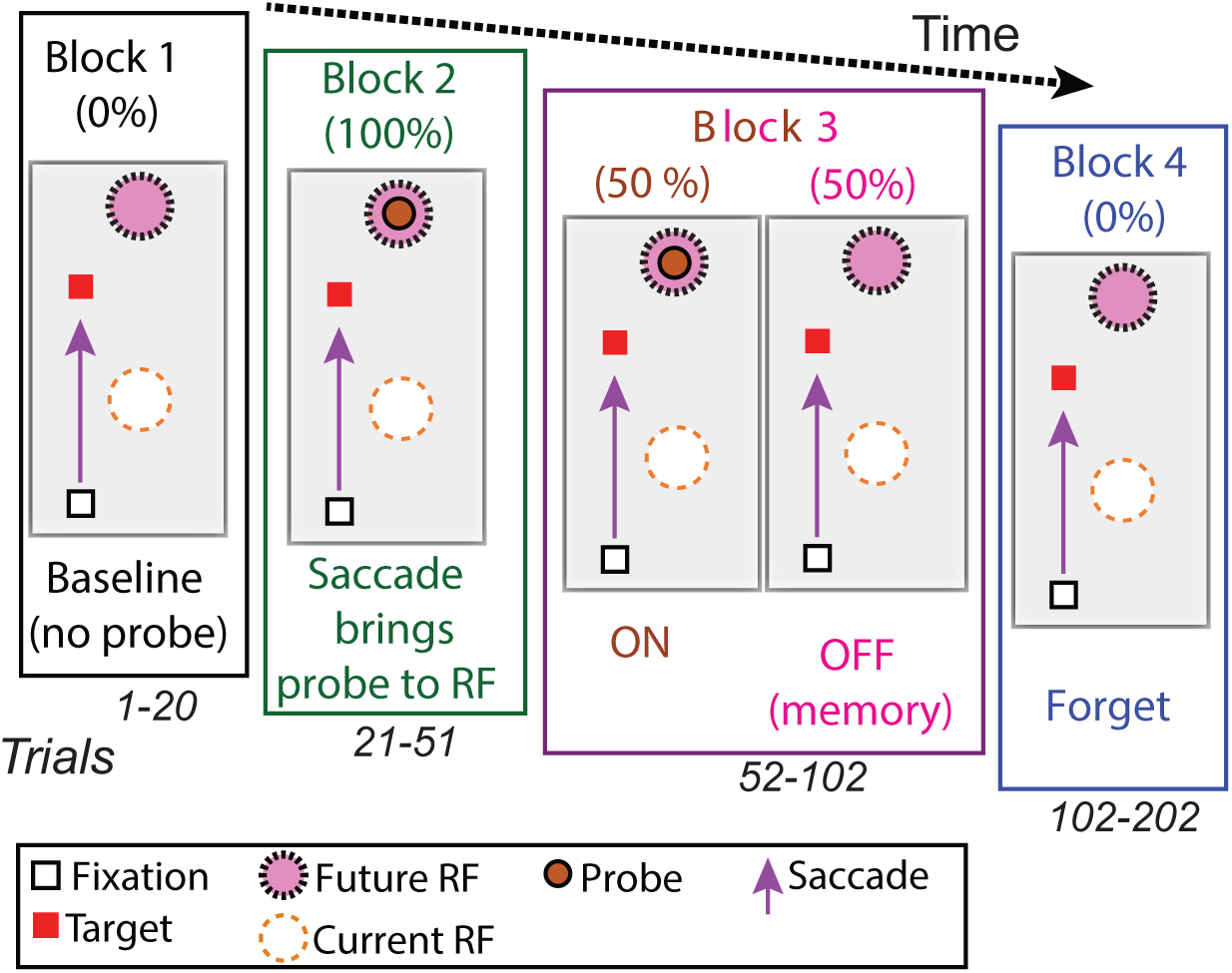
The basic memory task. The different trials types were performed in blocks. In Block 1, the monkey made a visually-guided saccade to a point outside the neuron’s response field, to show that the saccade itself does not evoke neural activity. In Block 2, the monkey made the same saccade, but now the saccade brings the probe stimulus into the receptive field. If the monkey made a saccade to the probe stimulus, the trial was terminated. In Block 3, for half of the trials, the probe stimulus appeared on the screen and was brought into the RF by the saccade. These trials were pseudorandomly intermixed with trials in which no probe stimulus appeared and the monkey made the same saccade. In block 4, the monkey made the same saccade but, as in Block 1, the probe stimulus never appeared.

We defined environmental memory the following three criteria: 1.) A significant difference (p < 0.0001 by Wilcoxon Ranksum test) in the post saccadic response in Block 1 and the probe stimulus - absent trials in Block 3 (Figure 3, black traces and raster symbols versus magenta traces and raster symbols). 2.) No significant post-saccadic response in Block 1 (Figure 3, black traces and raster symbols). 3.) A response to the probe stimulus in Block 2 (Figure 3, green traces and raster symbols). Of note, the visual response to the probe stimulus often occurred with a decreased latency (compared to the visual latency of the cell) typical of perisaccadic remapping (Umeno and Goldberg, 1997). Neurons responded similarly in Block 3 on the half of trials in which the probe stimulus appeared in the future receptive field (Figure 3, red traces and raster symbols). However, on the interleaved half of the trials in Block 3 in which a probe stimulus did not appear, the cells responded when the monkey made a saccade bringing the vanished probe location into the future receptive field (Figure 3, magenta traces and raster symbols). This response, which occurred after the saccade brought the spatial location of the probe stimulus into the receptive even though no probe appeared on that trial, is the environmental memory response. Finally, after a block of forgetting trials in which the probe stimulus never appeared (Block 4 trials, identical to Block 1 trials), the environmental memory response waned and, in some cases, disappeared entirely as in the example from monkey A (Figure 3, left panel).

**Figure 3.**
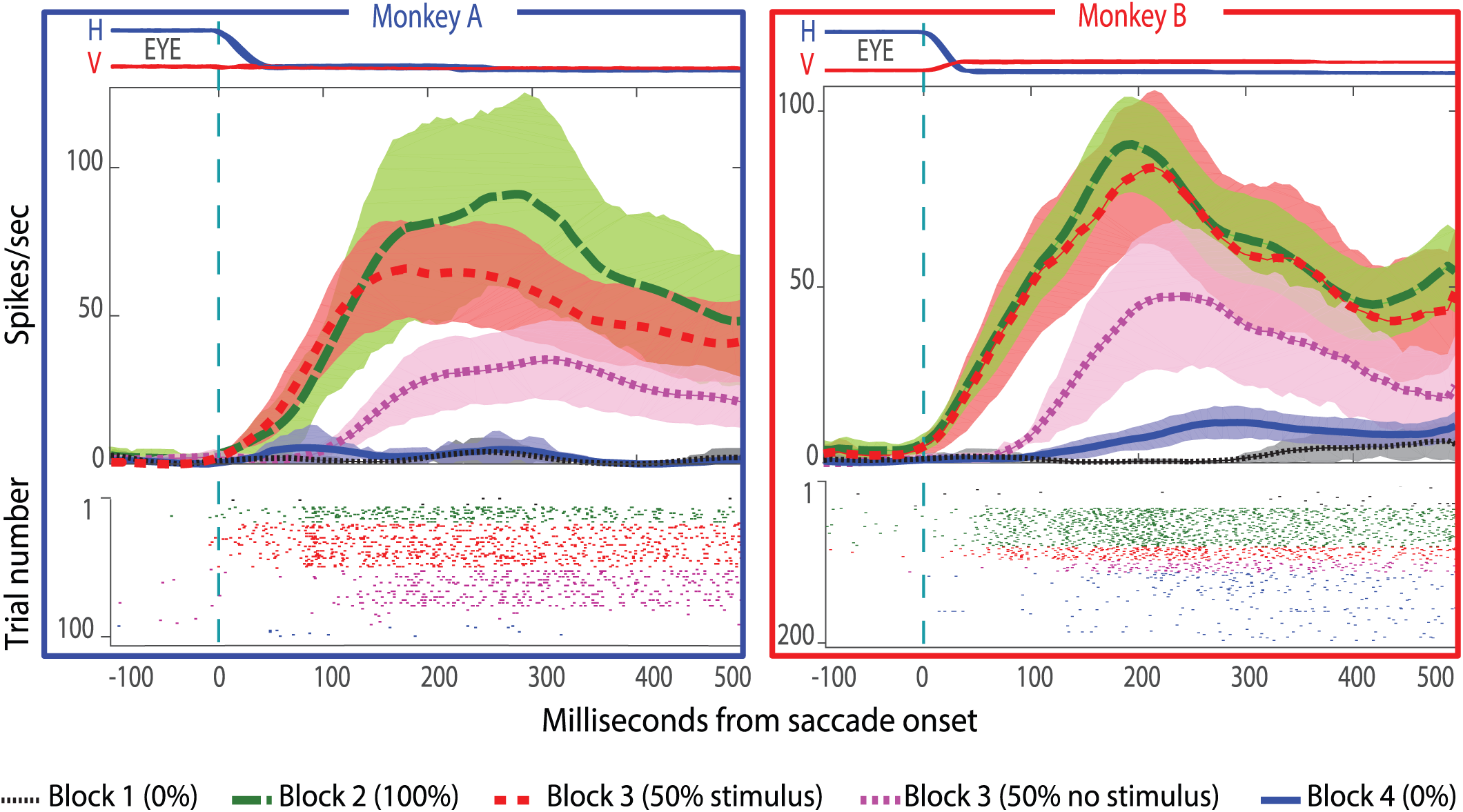
Single cell responses in the basic memory task from Monkey A (left panel) and monkey B (right panel). Neural activity aligned to the beginning of the saccade. Shaded regions in the peristimulus time histograms (PSTHs) are standard errors of means calculated using all the trials in the given block. Black traces and raster dots: activity in Block 1, stimulus absent. Green traces and raster dots: activity in Block 2, 100% stimulus present. Red traces and raster dots: activity in stimulus-present trials in Block 3. Magenta traces and raster dots: activity in probe stimulus-absent trials in Block 3. Here, there is a brisk response that shows the expected latency advance evoked by predictive remapping (Duhamel et al., 1992). Note that the latency of the memory response is nearly 100 ms longer after the saccade than the predictive response. Blue traces and raster dots: activity in Block 4, stimulus absent. The memory decays after many (up to 100) trials. H and V are horizontal & vertical eye movements smoothed using a 10 ms sliding causal filter.

The single cell results in Figure 2 were verified by the larger samples. For monkey A, we studied all 152 visually-responsive neurons which responded with delay or postsaccadic activity in the visually-guided delayed saccade task, of which 29 (20%) exhibited environmental memory, as previously defined. For monkey B, we studied 14 similar neurons, of which 9 (64%) showed significant environmental memory responses (p < 0.0001 by Wilcoxon Sign Rank) (Figure 4a, Monkey A in blue and Monkey B in red). The environmental memory response was weaker than the visual response evoked when the monkey made a saccade that brought the stimulus into the receptive field (Figure 4b, p < 0.001.)

**Figure 4.**
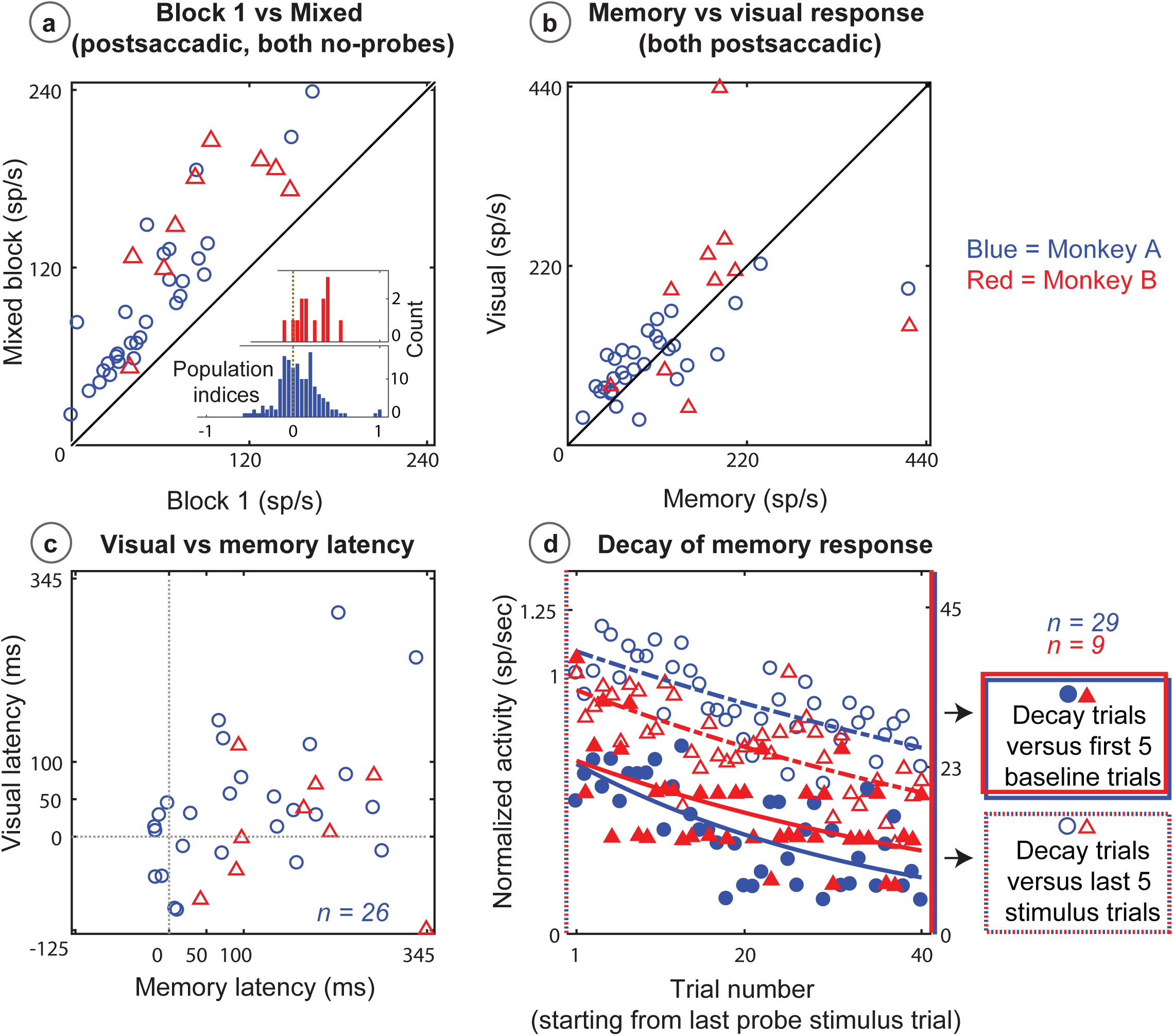
Population data for the basic memory task for units which showed significant differences between baseline and memory blocks (trial per trial peak comparisons, Wilcoxon ranksum, p<0.0001). **A.** Comparison of peak activity in the baseline and memory conditions (Block 1 vs Block 3, no probe stimulus trials). Each point represents a single cell. Cells with memory activity lie above the x=y line (diagonal). Monkey A: blue circles; Monkey B: red triangles. Only significant cells are shown. Insets: index (Block 3 versus Block 1) histograms for the whole population, out of which only statistically significant units were selected for the scatter plot. A ranksum and KS test confirmed memory activity as they both showed that the index distribution for the parent population was positively skewed. **B.** Peak memory activity plotted against visual activity. **C.** Latency of memory response versus the visual response for cells. **D.** Normalized decay activity. Memory response gradually decays after the last trial in which the probe appeared. The regression lines are data fits using a first order exponential function. As indicated in “D” legend, open circles and triangles are neural activities during decay trials compared to (normalized by) the mean of the last 5 trials with the probe stimulus (left y-axis). Shaded circles and triangles are decays compared to the mean of the first 5 baseline trials (right y-axis).

For all recorded cells, the population indices were significantly skewed towards reflecting memory activity. To investigate the population median shift, we used a non-parametric and robust skewness estimator called the medcouple measure (Brys et al., 2004) (abbreviated “medc” henceforth). Unlike classical histogram descriptors such as skewness and kurtosis, this approach finds the scaled median difference between the left and right sides of a distribution. Its values range from -1 to 1 (left to right skewed, respectively). Using this analysis method for Block 3 versus Block 1 indices (Figure 4a inserts) revealed that the population data was skewed towards a memory response (positive medc, 0.6 for monkey A, and 1 for monkey B). We also compared the index histograms with a mean-deducted version of the same distribution and found that memory activity significantly increased the median (ranksum p < 0.05).

In general, the latency of the environmental memory response was greater than the visual response which occurred when a saccade brought the stimulus into the receptive field (Figure 4b). Some, but not all, of the cells exhibiting environmental memory had a shorter visual latency than one would expect given the minimal latency to a stimulus flashed in the receptive field. This latency advance is characteristic of the remapping response (Duhamel et al., 1991). Of note, however, a number of cells exhibiting environmental memory did not show the remapping latency advance. In addition, because the memory responses for some neurons increased gradually, we were unable to establish memory latencies for all neurons using our latency method.

During Block 4, in which the probe stimulus no longer appeared, the memory response gradually decayed. On an individual cell basis, this decay (measured in the memory window as described in the methods section) was quite noisy and variable among cells. As a population, the normalized decay activity could be fit to a first order exponential with remarkably similar decay rate for each monkey irrespective of tasks used (Figure 4d). To control for fast memory buildups and/or in-between block fluctuations, we normalized the (compared) decay activity with Block 1 (Figure 4d, left y-axis) and also stimulus presentation block (right y-axis). For the basic memory task, monkey A had a decay rate constant of 0.13/0.33 trials (i.e. versus stimulus/block one trials) (R^2^ = 0.45/0.40). Monkey B had similar results (decay rate constant: 0.16/0.22; R^2^ = 0.48/0.23). For both monkeys and comparisons, the first and last 10 trials in the 40-trial forgetting block (Block 4) are statistically different (Wilcoxon ranksum, p < 0001).

#### Task 2: No-RF task

This task was designed to test whether environmental memory could be evoked in the absence of receptive field stimulation. The basic memory task demonstrated that LIP neurons respond when a saccade brings the spatial location of a previously presented (now vanished) probe stimulus into their receptive field. However, because we established memory by having the monkey make a saccade that brought a stimulus in into the receptive field, and then demonstrated the memory by having the monkey make the same saccade without a stimulus present, we could not know whether the memory response was independent of receptive field stimulation, or if it required visual stimulation to be established. Furthermore, we also could not determine whether the memory response was independent of the saccade used to establish it. To answer these questions, we used a modified version of the basic memory task, the No-RF task, in which the probe stimulus never appeared in the receptive field of the neuron (Figure 5). Trials in the first block of the No-RF task were identical to those in Block 1 of the basic memory task. In Block 1, the monkey made a visually-guided saccade (saccade 1) to a target outside of the neuron’s receptive field and no probe stimulus appeared (Figure 5, Block 1). In Block 2, two trial types were pseudorandomly interleaved: no-probe trials identical to those in Block 1 (requiring saccade 1), and trials in which the probe stimulus appeared, but a different saccade (saccade 2) was required. In the latter trials, the probe stimulus appeared in the location corresponding the neuron’s receptive field after saccade 1. However, instead of making saccade 1, monkeys were instructed to make a saccade 2, which relocated the neuron’s receptive field away from the probe stimulus location, thus preventing it from ever visually stimulating the cell. Block 3 was identical to Block 1, allowing us to measure the decay rate of the memory response. In Block 4, the monkey was instructed to make saccade 1 and the probe stimulus was presented in the receptive field, visually stimulating the cell. This enabled us to compare the memory and visual responses, and ensure that we had not lost the neuron during Block 3 while we observed the memory response decay.

**Figure 5.**
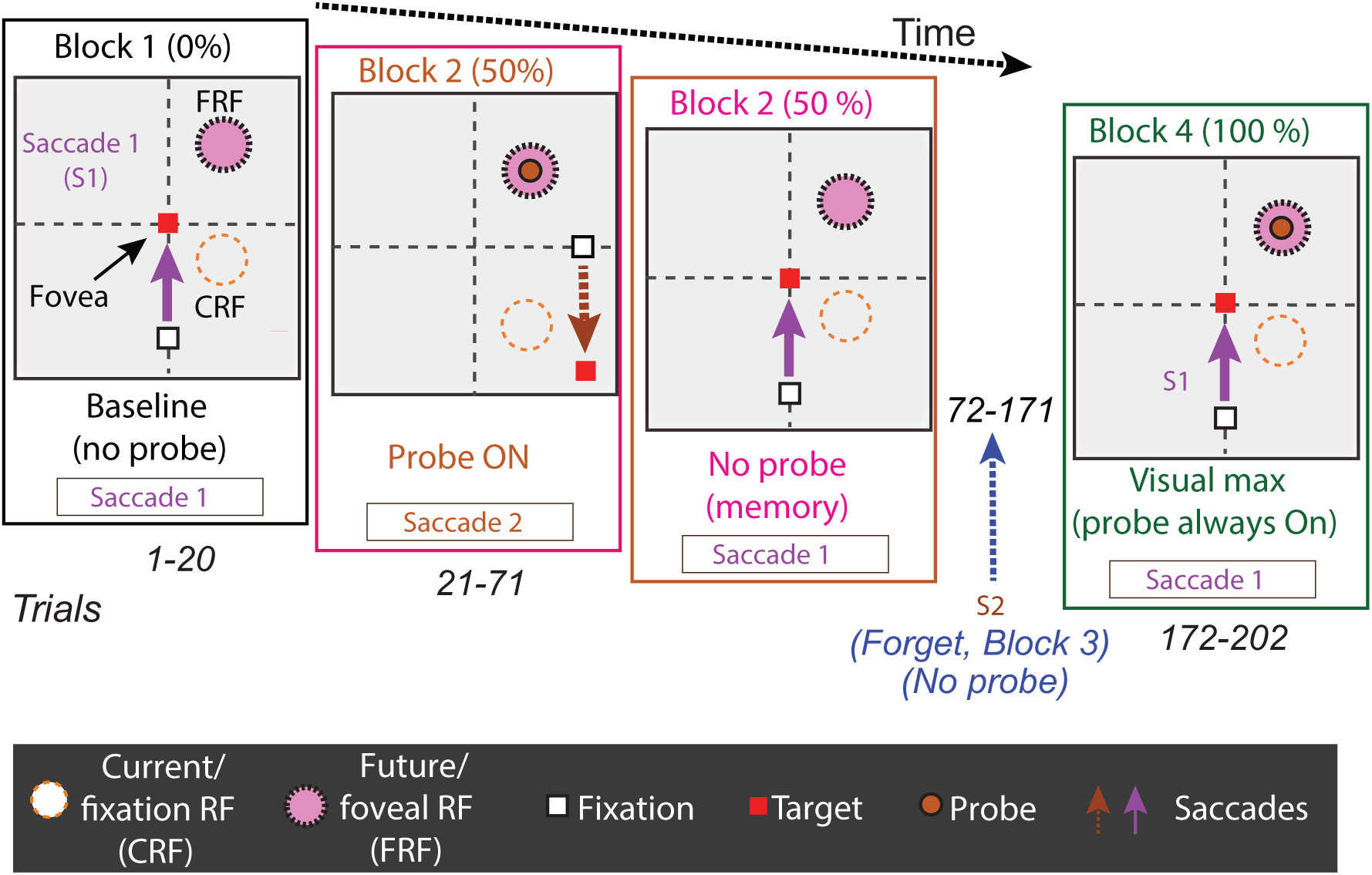
The No-RF single-saccade task. Trials occurred blocks. The first block was used to establish the baseline activity of the neuron for saccade 1, prior to introduction to the probe stimulus. Block 2 introduced a task-irrelevant probe stimulus in half of the trials. In the half of trials where the probe stimulus appeared, the monkey was instructed to make a different saccade (saccade 2), which relocated the cell’s receptive field away from the probe location (Block 2, orange). In the other half of trials, no probe stimulus was presented and the monkey made saccade 1, bringing the location of the (previously presented) probe stimulus into the cell’s receptive field (Block 2, magenta). In Block 3 (not shown, identical to Block 1), to measure the decay of the memory response, the monkey was instructed to make saccade 1 and no probe was presented. Finally, in Block 4, to measure the visual response of the cell to the probe, the monkey was instructed to make saccade 1, and the probe stimulus was presented in the cell’s receptive field.

We studied 169 neurons (139 in monkey A, 30 in monkey B) in the no-RF task. Of these, 29 (17%) LIP neurons showed environmental memory (as previously defined). The environmental memory response was observed (Figure 7a) despite the facts that 1) the probe stimulus never appeared in the neuron’s receptive field, 2) the saccade used to establish the memory response (saccade 2) differed from the saccade used to evoke the memory response (saccade 1). Finally, to obtain a visual response to the probe stimulus for comparison with the memory response, in Block 4 we placed the probe stimulus at the location corresponding to the cell’s receptive field after saccade 1 and instructed the monkey to make saccade 1. We found that the probe stimulus drove the cell robustly (Figure 5, block 4), confirming the geometry of stimulus placement.

Similar to the basic memory task, for the No-RF task population indices (Block 3 versus Block 1, Figure 7a inserts) were shifted to positive median values (medc: 0.5 for monkey A, and 1 for monkey B; versus-mean-deducted ranksum p < 0.03). Also similarly, the memory response was generally less robust (Figure 7b) and had a longer latency than the neuron’s visual response (Figure 7c). As with data from the basic memory task, the normalized decay activity of the population (Figure 7d) could be fit to a first order exponential (monkey A had a decay rate constant of 0.03/0.10; R^2^ = 0.49/0.57; Monkey B: decay rate constant of 0.08/0.10 trials; R^2^ = 0.02/0.05;). Significant p-values were found using a Wilcoxon test between trial mean comparisons of the first and the last 10 trials in the 40-trial decay epoch considered (p < 0.03, for both monkeys and comparisons).

#### Task 3: Two-saccade task

LIP plays an important role in choosing targets for visual and memory-guided saccades (Gnadt and Andersen, 1988, Wardak et al., 2002, Ipata et al., 2006). Because the plan to make a saccade into a cell’s receptive field is reflected in LIP with an increase in activity corresponding to that location, it is essential to rule of the possibility that the memory signal is actually a saccade plan. To address this, we trained Monkey A in a two-saccade task. In this task, the monkey made an initial saccade that brought the spatial location of the (vanished) probe stimulus into the receptive field, followed by a saccade which was directed away from the probe location (away from the receptive field) (Figure 6).

**Figure 6.**
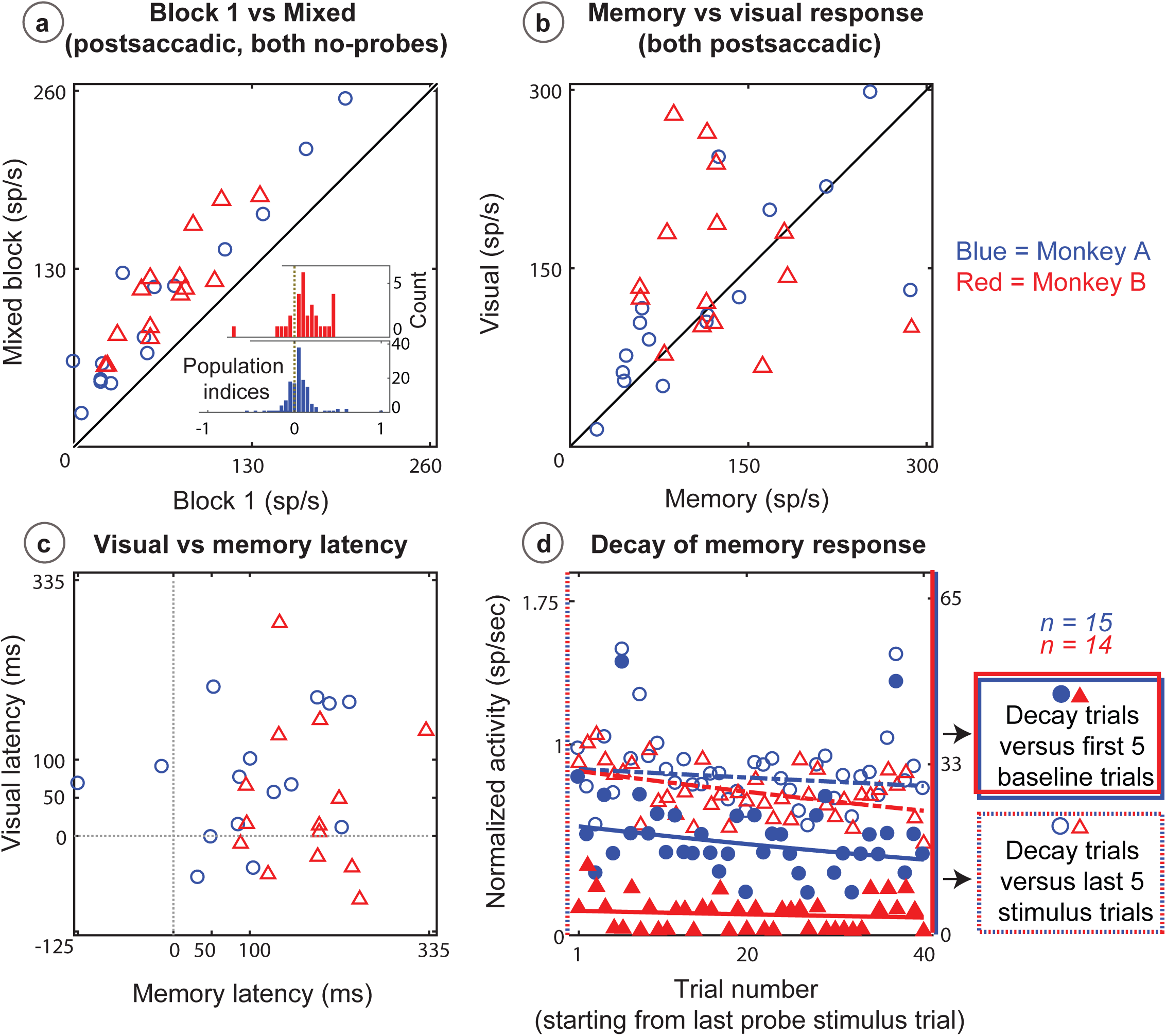
The No-RF two-saccade task. The block design is the same as the no-RF single-saccade task except the monkey was required to make two saccades. The requirement of a second saccade, away from the cell’s receptive field, prohibits the planning of a saccade to the probe stimulus location. Block 3 was identical to Block 1 (not shown), and was used to measure the decay of the memory signal.

In this task, we studied 157 neurons from monkey A; 11% (i.e. 18/157) of which exhibited a statistically significant environmental memory. The memory signal was evoked even though the monkey was planning a saccade away from the receptive field, and would in fact would be punished (by receiving no reward) should it make the saccade to the probe location (Figure 8a). Block 3 versus Block 1 indices for the population (Figure 7a insert) behaved the same as in the other two tasks, in that the median memory response in Block 3 was higher than the baseline response in Block 1 (medc: 0.4; versus-mean-deducted ranksum p < 0.05). The strength of the memory response was not as great as the visual response (Figure 8b), and the latency of the memory response was longer than that of the visual response (Figure 8c). Compared to the other tasks using the same monkey, decay activity in this task resulted in a similar population response (Figure 8d). The decay constant was 0.12/0.15 trials, with R^2^ of 0.09/0.19. A Wilcoxon test between the first and last 10 trials in the 40-trial epoch showed significant decay (p < 0.01, both comparisons).

**Figure 7.**
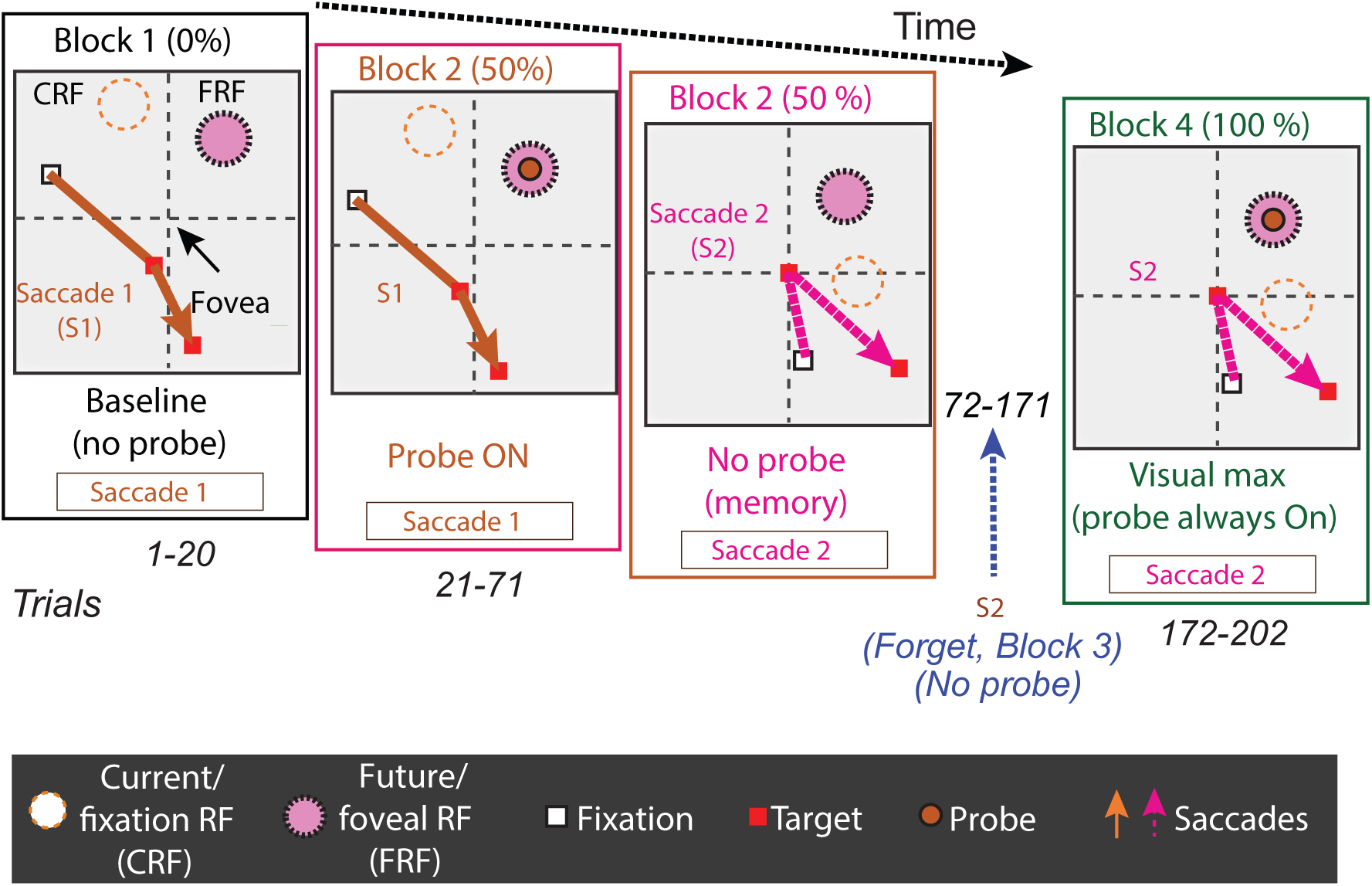
Population data for the no-RF task for units which showed significant differences between baseline and memory blocks (trial per trial peak response comparisons, Wilcoxon ranksum, p<0.0001). Color codes and statistical analysis are as in Figure 4. A. Comparison of peak activity in the baseline and memory blocks of trials. Each point represents a cell. Cells with memory activity lie above the x=y line (diagonal). Monkey A: blue circles; Monkey B: red triangles. Only significant cells are plotted. B. Peak memory activity plotted against visual activity. C. Latency of memory response versus the visual response. D. Normalized decay activity. Memory response gradually decays after the last trial in which the probe appeared. The regression lines are data fits using a first order exponential function.

**Figure 8.**
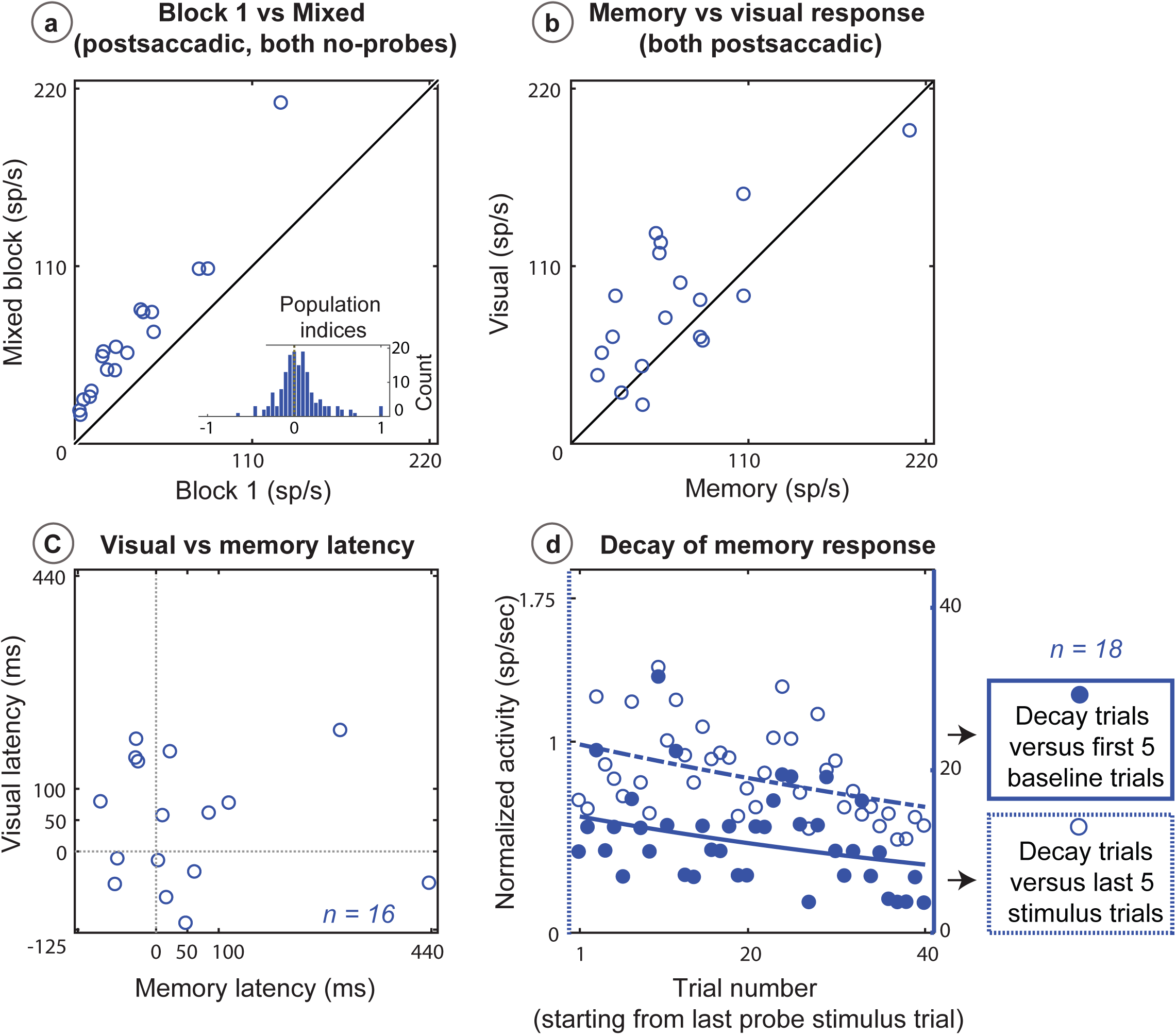
Population data for the two-saccade task for units which showed significant differences between baseline and memory blocks (trial per trial peak response comparisons, Wilcoxon ranksum, p<0.0001). **A**. Comparison of peak activity in the baseline and memory blocks of trials. Each point represents a cell. Cells with memory activity lie above the x=y line (diagonal). Data from Monkey A only. Only significant cells are shown. **B.** Peak memory activity plotted against visual activity. **C.** Latency of memory response versus the visual response. **D.** Normalized decay activity. Memory response gradually decays after the last trial in which the probe appeared. The regression line is a data fit using a first order exponential function.

## Discussion

In this set of experiments, we demonstrate that LIP neurons exhibit an environmental memory signal, which lasts over a duration of multiple trials, decays slowly, and updates with eye movements. In addition, we demonstrate that a neuron does not require visual stimulation or a particular training saccade to establish memory activity. A similar signal was found in the frontal eye field (Umeno and Goldberg, 2001), in visual and visuomovement cells, but not movement cells. These preliminary experiments, however, had two confounds. First, the memory was always established by stimulating the receptive field. Second, the saccade that evoked the memory was identical to the saccade used to establish it, so the memory could have been related to the saccadic process. In our experiments, we show that memory can be evoked by a stimulus that never appeared in the neuron’s receptive field, and that the memory response manifested even when the saccade used to evoke it differed from the saccade used to establish it. Furthermore, as in the FEF (Umeno and Goldberg, 2001), memory activity occurred both in neurons which did exhibit saccadic remapping as well as those that did not. In sum, these results suggest that LIP has access to a craniotopic representation of the visual world despite the fact that all of its explicit responses are retinotopic, and that this memory response cannot be simply ascribed to saccadic remapping or saccade planning.

Our results are consistent with previous observations regarding the role of parietal cortex in spatial memory. Patients with bilateral parietal lesions cannot point to an object in their room with their eyes closed, although they can easily do it with their eyes open (Farah et al., 1988). This suggests that these patients do not have access to a remembered spatial representation of their environment. Paradoxically, verbally reporting this memory requires conversion to a retinotopic frame. This was best shown in the classic paper by Bisiach and Luzzatti (Bisiach and Luzzatti, 1978), in which two Milanese patients with right parietal damage were asked describe their memory of the Piazza del Duomo in Milan. When they described it as if they were standing with their back were to the cathedral they remembered only the landmarks to the left of the cathedral. When they described it as if they were facing the cathedral they only remembered the landmarks on the other side of the square. This experiment demonstrates two key ideas; 1. The long-term memory of environments is stored in supraretinal coordinates in an area unaffected by a parietal lesion. 2. Retrieving that memory goes through a retinotopic process that requires the parietal cortex. We suggest that LIP is a part of the network that transforms a supraretinal memory into a retinotopic signal that is useful for the motor system.

This raises the question of how a supraretinal memory can be established. There are two known mechanisms by which LIP can create a spatially accurate representation of space despite a constantly moving eye. The first is through remapping, in which a neuron will respond immediately after a saccade brings the spatial location of a recently-vanished stimulus into its receptive field (Duhamel et al., 1991, Sun and Goldberg, 2016). This effectively changes the spatial origin of the retinotopic coordinate system from the presaccadic fixation point to the saccade target. The next saccade could just as easily remap the current retinotopic representation into a coordinate system centered on the next saccade target, and so on through a number of saccades. This is unlikely, however, because many cells that show environmental memory do not show pre-saccadic remapping when a saccade brings a stimulus into its receptive field.

The second mechanism creates a spatially accurate retinotopic representation by developing a retinotopic representation from a craniotopic representation. There is excellent human psychophysical evidence that the saccadic system has access to a craniotopic representation of space. Karn et al (Karn et al., 1997) showed that there is little difference in the inaccuracy of memory-guided saccades after two or six intervening visually-guided saccades. Reliance on a remapping mechanism for accuracy would entail the concatenation of errors from saccade to saccade, whereas access to a craniotopic coordinate frame would enable a more stable representation of space across multiple saccades. In a similar experiment, Poletti et al (Poletti et al., 2013) asked humans to make different numbers of intervening saccades between the presentation of a target for a memory-guided saccade and the actual saccade. For the first few intervening saccades, the variability of the memory-guided saccade increased linearly, as expected if the error of each remapping process would concatenate. However, after the first few saccades, the increase in variability decreased with each subsequent saccade, as if the representation of the saccade target had shifted to a craniotopic frame which would not increase its error with each saccade. To explain this phenomenon, they proposed a model which entails a shift from an early remapping mechanisms to a subsequent craniotopic mechanism. This late craniotopic representation could arise from a gain-field mechanism. Visual responses in LIP are linearly modulated by the position of the eye in the orbit, the gain field. The eye position signal that modulates the gain fields in LIP could come from the representation of eye position in area 3a (Zhang et al., 2008). It is mathematically trivial to calculate target position in craniotopic coordinates from retinotopic gain fields (Zipser and Andersen, 1988, Salinas and Abbott, 1997, Pouget and Sejnowski, 1994). Although gain fields are inaccurate immediately after a saccade (Xu et al., 2012), they could contribute to a late, accurate, craniotopic representation that develops when the eye position signal is accurate, as one would expect if the eye position signal arose from proprioception. The neurons that we studied did not have eye position modulation of their visual responses – such neurons often have a phasic and tonic postsaccadic response that arises from the eye position signal itself (ref), although the craniotopic representation to which they must have access may arise from a gain field calculation involving other LIP neurons.

Data from patients with parietal lesions, such as the Bisiach-Luzzatti result previously described, suggest that although LIP might calculate target position in craniotopic coordinates, the parietal cortex does not store the results of the calculation. Instead, the representation of space could be stored elsewhere, possibly in the medial temporal lobe. O’Keefe discovered that neurons in the rat hippocampus (the ‘place cells’) discharge when rats enter spatial location that they have seen before. A few studies have duplicated this result in the monkey hippocampus (Ono et al., 1993, Rolls et al., 1989). Grid cells in entorhinal cortex are thought to be the precursors of hippocampal place cells (Moser et al., 2008) and these have been found in the monkey (Buffalo, 2015) as well as in humans (Jacobs et al., 2013). It is known that there is a direct, reciprocal connection between the parahippocampal gyrus and LIP (Suzuki and Amaral, 1994, Baizer et al., 1991). Thus, LIP could send an eye-position modulated visual signal to the medial temporal lobe, which could serve as the building block for a craniotopic memory. The medial temporal lobe could then use this activity to generate the craniotopic component of place cells and send this craniotopic signal back to LIP. LIP itself would only be activated when the spatial location of the vanished stimulus can be described by a retinotopic vector from the center of gaze to the receptive field of the neuron. Thus, LIP has access to a craniotopic representation, but expresses only a retinotopic representation.

Although early models posited that the oculomotor system uses a craniotopic representation (Zee et al., 1976), it is clear that the parietal cortex sends a spatially accurate retinotopic signal to the oculomotor (Pare and Wurtz, 1997) and skeletomotor (Batista et al., 1999) systems. Even auditory stimuli, which are initially coded in craniotopic coordinates, are converted to retinotopic coordinates for the oculomotor (Jay and Sparks, 1987b, Jay and Sparks, 1987a) and skeletal motor system (Grunewald et al., 1999) (Cohen and Andersen, 2000) in parietal cortex, and the oculomotor system of the superior colliculus (Jay and Sparks, 1984). We suggest that the environmental memory signal in LIP is an example of the conversion of a craniotopic to a retinotopic signal for memory and spatial attention as well as for motor control

## Methods

The Animal Care and Use Committees at Columbia University and the New York State Psychiatric Institute approved all of the animal protocols in this study as complying with the guidelines established in the United States Public Health Service Guide for the Care and Use of Laboratory Animals. We used one female and two male rhesus monkeys (Macaca mulatta) with weights between 6 and 13 kg.

Monkeys were first trained to sit in a primate chair using a pole and collar technique. After chair training, monkeys were surgically fitted with acrylic implants, anchored with titanium or ceramic screws, outfitted with a headpost which stabilized the monkeys head while sitting in a primate chair. During the same surgery, scleral search coils were implanted around both eyes (Judge et al., 1980) and the coil wire brought out to a plug on the acrylic. After basic behavioral training, a craniotomy was performed to expose the intraparietal sulcus (using coordinates determined from a structural MRI), over which a recording chamber was affixed in in the acrylic.

MRI’s were obtained after tranquilizing the monkey with ketamine and atropine for transport to the MRI lab. The monkey was then anesthetized with endotracheal isoflurane, and positioned in a Kopf MRI compatible stereotaxic instrument. To create the images, we used a custom-made rigid and flexible RF array coils using GE MR750 and a GE 1.5-T Signa scanners. We used custom Matlab software, Osirix, GE’s MediaViewer, and DicomWorks to analyze the data.

We performed all surgeries using ketamine/isoflurane general anesthesia using aseptic surgical technique. The monkey recovered fully before testing restarted. During testing, monkeys worked for their daily water intake and were supplemented with dried and fresh fruits. Monkeys’ weights and general health were monitored on every recording day, and at least once a week.

We controlled all experiments using the REX (downloadable from: https://nei.nih.gov/intramural/software) system (Hays et al., 1982) and recorded single-unit activity with glass-insulated tungsten electrodes introduced through a guide tube positioned plastic grid with 1mm spacing between possible penetrations (Crist et al., 1988). In general, recording sessions lasted between 5 and 8 hours depending on the stability of the cell and the monkey’s willingness to continue to work. We took care to avoid making penetrations with the electrode in adjacent grid holes from the previous day’s penetration in order to reduce tissue damage. We kept careful logs regarding the location and depth of each penetration an estimate describing cell types encountered, and the quality of the monkey’s behavior. During every testing and training session, we monitored monkeys using a closed-circuit camera and monitor.

We verified stimulus timing using a photoprobe which emitted a pulse for each event. We monitored the monkey’s eye movements using a CNC phase detector (Crist Instruments) to decode the search coil signal (Judge et al., 1980) and sampled the signal at 1 kHz. During testing, monkeys sat in a primate chair in a sound a sound-attenuated Faraday room. We performed these experiments in two experimental setups. In one set up, the monkeys sat 57 cm away from a CRT monitor (ViewSonic Professional Series P225F) with a refresh rate of 120 Hz. In the second set up, stimuli were presented with a Hitachi CPX275 LCD projector with a refresh rate of 60Hz, on a screen 72 cm from the monkey.

Once we isolated an LIP neuron and confirmed that it displayed canonical visual, delay and perisaccadic responses, we probed the boundaries of LIP neuron receptive fields by having monkeys make memory-guided saccades (Hikosaka and Wurtz, 1983) to targets appearing in locations which were inside and outside of the cell’s receptive field. In some cases, we used a fixation task to characterize its receptive field by having the monkey fixate a point while stimuli briefly flashed (50ms) pseudo-randomly on the screen, generating a map of locations that evoked visual responses (measured 50-150 ms after the flash). We ensured that saccade target and probe stimuli locations were positioned correctly with respect to future and current receptive field locations to avoid unintentional contamination by a visual response.

We transformed the REX data into a form analyzable by MATLAB (MathWorks, Natick, MA) using the REXTOOLS software (downloadable from https://nei.nih.gov/intramural/software). Although the REX system provides an estimate of activity using the MEX (downloadable from https://nei.nih.gov/intramural/software) spike sorter, for the analyses in this paper we used the MEX system to digitize the neural data and sorted the spikes offline using the MEX offline analysis program. The custom codes, and raw data from every cell analyzed in this paper are provided as .m, and .csv files.

Before performing detailed statistical analysis, we first investigated if the data came from standard normal distributions, using two-sample Kolmogorov-Smirnov test. When we tested the mean neural activity in the response windows (intervals described below), all of our data sets were found to be distributed non-normally (non-parametric).

To show that the statistically significant cells in all three experiments were not just one side of a symmetric distribution we calculated a memory index to quantify the degree to which each cell manifested memory activity, comparing postsaccadic activity in Block 1 (R_pre_) with postsaccadic activity in Block 3 (R_mem_):

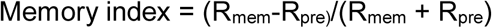

All index distributions were not normally distributed (by KS test), and we showed that the population median was significantly different from zero by using a Mann-Whitney ranksum test to compare each sample with a test sample identical to the measured sample with the measured mean subtracted from each value. In addition, we calculated medcouple, a non-parametric histogram skewness measure (Brys et al., 2004). This tool is not affected by unsymmetrical tails and outliers, and is also not based on classical skewness third moment. Since this measure doesn’t depend on mean or standard deviation, it gives an accurate description of a skewed distribution. Its results are bound between values: -1 (left skewed), 0 (symmetric), and 1 (right skewed).

All cells reported here had both statistically significant visual and memory responses (Mann-Whitney ranksum p-value for median activity between baseline and response window < 0.05). We have excluded units with memory activity if median activity in Block 1 (baseline recording block) pre- and post-saccadic was statistically different. In other words, even if we could find significant memory activity well beyond what can be described by either eye position or saccadic activity effect, we are not presenting them here.

To quantify the magnitude of the memory and visual responses, we did a comparison of the peri-stimulus time histograms (PSTHs) during baseline, visual and memory trials, for identical epochs aligned on saccade onset. PSTHs were constructed from 2 ms spike bins, smoothed using a 10 ms sliding causal filter. Baseline calculations were made in the –426 to –226 ms (pre-saccade onset) window. To quantify the decay, perisaccadic memory and visual responses, we selected the time point of peak activity in the window from 125 ms before the saccade onset (to include any predictive remapping) to 500 ms after. We then looked at the median of raw spike count over a window extending 100 ms before and after this point to calculate response latencies, as described next.

To obtain a latency value of the memory response, we used a median threshold-crossing and sustained response method. In short, this analysis compared the median activity in the baseline and response windows. A “cut off” value was derived by determining the spike count values which would fall above at least the 75% percentile of the baseline median. By comparing the median spike counts in the response window to this “cut off” value, a latency time could be determined when a minimum of five bins (of 2 ms each) contained a greater spike count than the “cut off” value. The first of these bins was taken as the response latency, and verified visually by looking at the corresponding time on the PSTH. Since the baseline activity can vary between different trials in the same experiment, we first chose a window within the time from fixation to -226 ms before saccade onset. Within this wide window, we made 200 ms wide sliding windows, in 1 ms steps. By finding the median spike count within these smaller windows, we picked the window closest to the start of the response window (but still have the same median as the population of these several 200 ms wide windows). This median was chosen to look at the response window in which a threshold crossing for at least 20 ms was picked as the response latency.

We could not calculate latencies for a few neurons, because the baseline (pre-saccadic) memory block response was as high as post-saccadic activity. There were also a few other units whose latencies were automatically picked by the aforementioned median method but a more precise time point could be picked by doing yet another median-based analysis. For these units, we implemented Wilcoxon ranksum comparison with the baseline median and a sliding 50 ms window, stepped 1 ms, in the response epoch. The first such 20 ms window, where there was a statistical difference was observed was picked as the latency point.

We observed that the memory response continued to manifest for almost up to 100 trials after the last probe stimulus trial, and so we analyzed the “forgetting” or “decay trials”. While individual cell results were noisy and there were differences between cells, we normalized each cell’s decay activity to its peak memory activity and fit the data to a first or second order polynomial. We fit the population data similarly.

## Acknowledgments

We thank, Moshe Shalev and Girma Asfaw for veterinary care, Yana Pavlova and Vincent Sanchez for technical assistance, John Caban and Matthew Hasday for machining, Glen Duncan for electronic and computer assistance, and Latoya Palmer, Cherise Washington, and Lisa Kennelly for facilitating everything. This research was supported, in part, by grants from the Keck, Gatsby, Kavli, Zegar, and Dana Foundations and the National Eye Institute (R24 EY-015634, R21 EY-017938, R21 EY-020631, R01 EY-017039, P30 EY-019007, and R01 EY-014978 to M. E. Goldberg, principal investigator; S. S. was also supported by training grant T32-EY-13933. M. S. was supported by NINDS training grant from 2T32MH015174-35, Brain & Behavior Research Foundation (NARSAD) 2013 Young Investigator Award.

## Author Contributions

S.S., M.E.G., and M.S designed the experiment; M.S., and S. S. collected the data, M.S. performed the analysis; M.S. and MEG wrote the paper with inputs from S. S.

